# Motivational trade-offs as evidence for sentience in bees: a critique

**DOI:** 10.1101/2025.04.04.647054

**Authors:** Jenny Read, Vivek Nityananda

**Author notes:** Corresponding authors: Jenny Read, Vivek Nityananda.

## Abstract

Establishing if insects feel pain can have far-reaching consequences for insect husbandry, commercial pollination and scientific research. Research in this field therefore requires careful experiments and strong evidence. One important criterion proposed for measuring insect sentience is the ability to show a motivational trade-off, in which “the negative value of a noxious or threatening stimulus is weighed (traded-off) against the positive value of an opportunity for reward”. A recent paper investigated motivational trade-offs in bumblebees and concluded that bees can trade-off heat against high sugar rewards. In this paper, we develop a signal detection model to highlight which features would be key to supporting the argument that motivational trade-offs are evidence of the capacity for experiencing pain. We then re-analyse the data from the original paper and find several limitations. Our own re-analysis finds no support for the final conclusions made by the paper. We therefore provide recommendations for future studies investigating the ability of insects to feel pain.

## Introduction

Establishing if insects feel pain can have far-reaching consequences for insect husbandry, commercial pollination and scientific research. Research in this field therefore requires careful experiments and strong evidence. Recent innovative approaches to investigate this question test whether insects fulfil a list of eight criteria, such as nociception, sensory integration, and flexible self-protection, mostly assessed behaviourally (1, 2). One important criterion is the ability to show a motivational trade-off, in which “the negative value of a noxious or threatening stimulus is weighed (traded-off) against the positive value of an opportunity for reward” (1). This criterion has, for example been used to investigate pain in other invertebrates like hermit crabs (3).

### Motivational trade-offs as evidence for sentience

For a motivational trade-off to be evidence of sentience, “enough flexibility must be shown to indicate centralized, integrative processing of information involving a common measure of value” (1). To see why, consider that even microbes could make *behavioural* trade-offs (e.g. (4)). They might, for example, move away from heat when the sugar concentration is uniform but move towards heat if there was a sufficient high sugar concentration (Figure 1A). However, this could occur even if information were combined only in the final chemical pathways onto the motor system, e.g. if an increase in sucrose concentration or a decrease in temperature causes the microbe’s flagellum to move it forward, while a decrease in sucrose or increase in temperature causes it to change direction randomly. The negative value of heat and the positive value of sugar would effectively be combined into a common measure of value (“S-H” in Figure 1), but this would be hard-wired through the interaction of chemotaxis and thermotaxis proteins with the flagellar motor. The microbe’s trading off of heat for sugar would therefore not be considered evidence of sentience.

**Figure 1.**
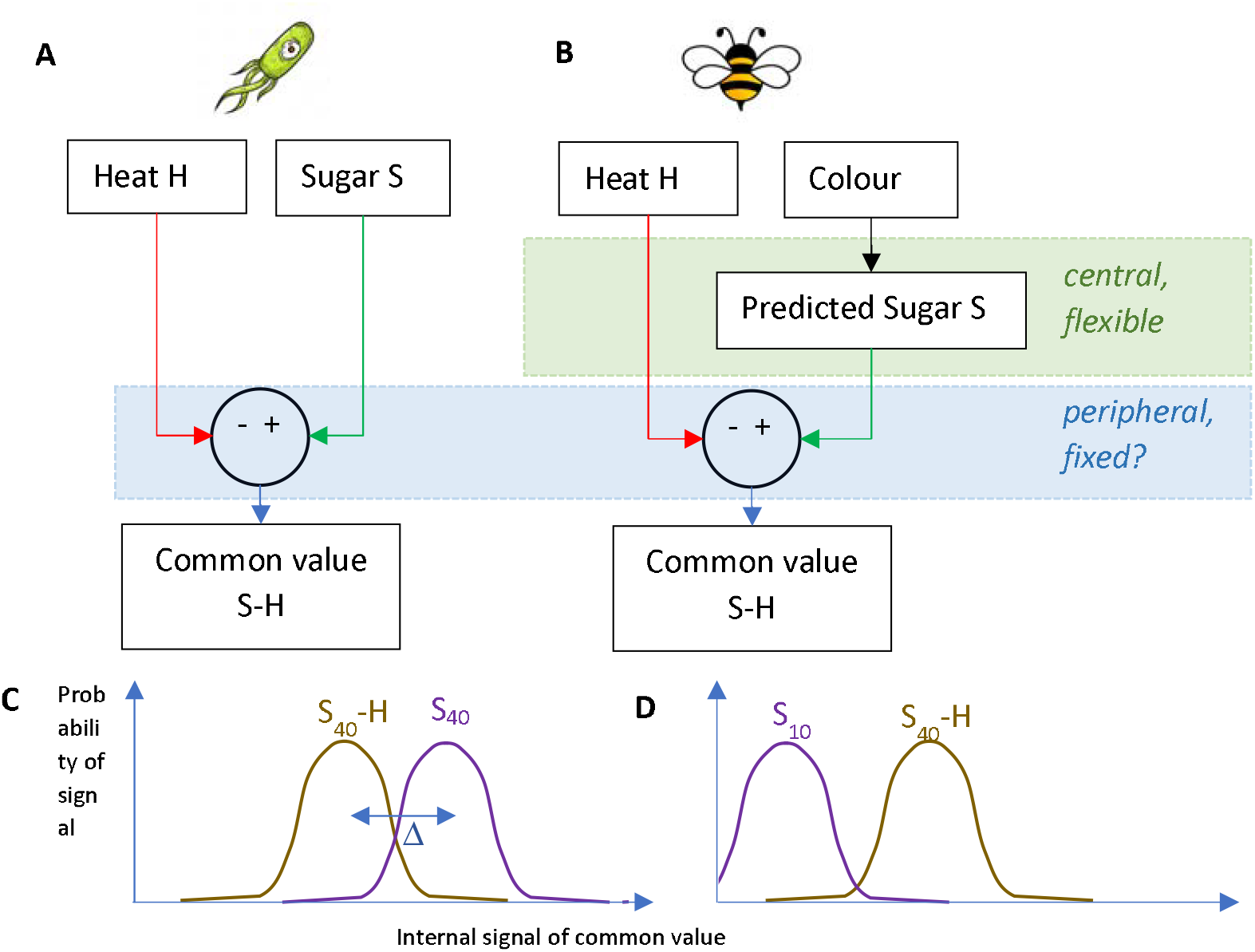
A: Even unicellular organisms can trade off stimuli of different value. For example, if sensed changes in heat and sugar both control flagellar rotation by the same ultimate molecular pathway, then both are effectively combined into a common value. B: In bees, behaviour can be controlled by an internal representation of predicted reward only recently learnt from an arbitrary mapping. This reward signal could be combined with signals about noxious stimuli to produce a common value signal that controls behaviour, but it is not yet clear whether the creation of common value is itself flexible or central. C, D: Simple signal detection theory account of how bees might decide between two options, based on their differing values. S_10_, S_40_ represent the reward value of unheated 10% and 40% sucrose respectively; H represents the aversive effect of heat. C: When both colours offer 40% sucrose but one is heated, bees are more likely to choose the unheated colour (purple) since this has the higher total value. D: When the 40% sucrose is heated but the other colour offers only 10% sucrose, bees reliably choose the heated colour (brown). We use Δ to represent the difference in value relative to the noise on the signal (Δ is often known as dprime in the literature).

A recent pioneering paper (5) investigated motivational trade-offs in bumblebees. The bees were first trained to associate high sucrose (40% solution) with one colour and another concentration (either 10%, 20%, 30% or 40%, one value per bee) with a different colour. The high-sucrose feeders were then heated. When both colours of feeder offered the same high-sucrose concentration, bees avoided feeding from the heated feeders. However, when the unheated colour offered low-sucrose, bees became relatively more likely to feed from the heated, high-sucrose feeders. This suggests they were trading off aversive heat for rewarding sucrose (Figure 1C and D, Figure 2).

**Figure 2.**
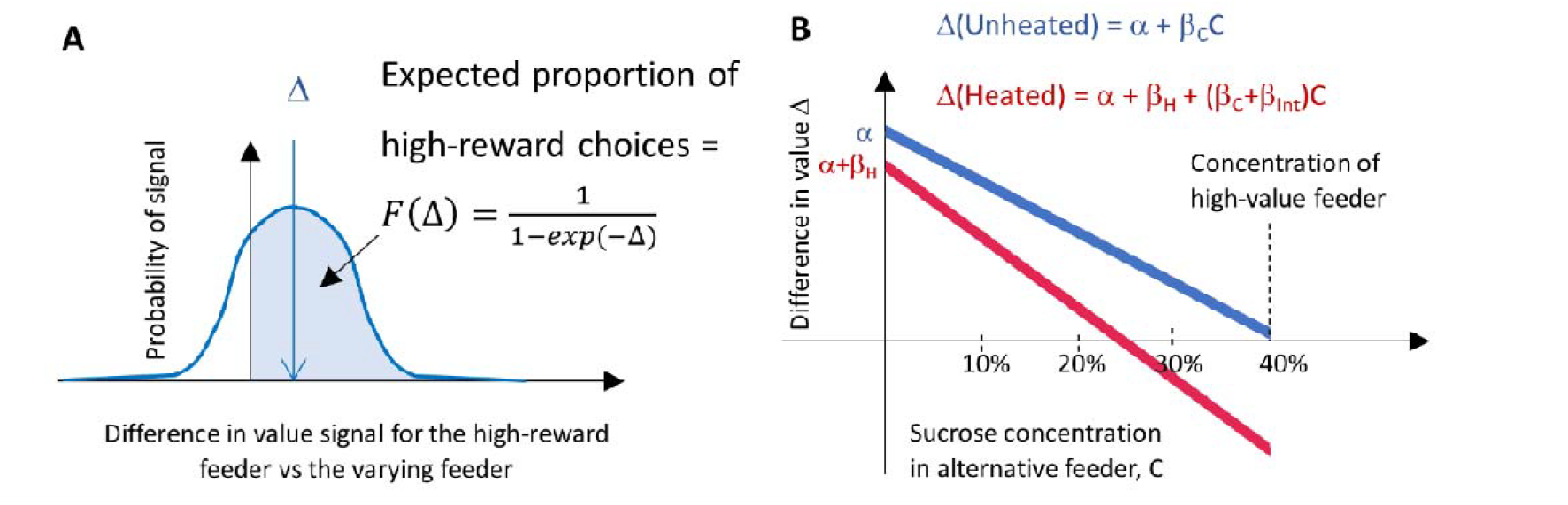
The signal detection theory implied by fitting a logistic function to this data. (A) The model assumes bees base their decisions on a noisy signal about the difference in value between the two choices (Figure 1C). The mean difference, relative to the standard deviation of the noise is represented by Δ. The GLMM models the proportion of high-value choices as a logistic function of Δ, corresponding to the shaded region of the distribution. (B) The GLMM models the difference in value Δ as a function of Concentration, C, and Temperature, T. α represents the model intercept, β_C_ the main effect of Concentration, β_H_ the main effect of Temperature, and β_int_ the interaction term.

This behaviour cannot be fully hard-wired, since the bees made choices based on the colour of the feeder without sensing the sucrose directly. The bees must have been generating their reward signal via an internal representation of the reward associated with each colour (Figure 1B). The authors conclude that “the trade-off [between heat avoidance against sucrose preference] is mediated in the central nervous system”.

This does not however necessarily follow from the data. In honeybees, neuronal activity correlated with the prediction of reward has been observed in the suboesophageal ganglion (6). The neuron involved projects to more central regions including the lateral protocerebrum and the mushroom bodies. But it doesn’t follow that the trade-off of predicted reward with nociception *also* occurs flexibly and centrally. It could, for example, occur peripherally through a fixed chemical pathway, as in the microbe example (Figure 1A, B). Thus, without evidence that the combination is indeed central and flexible, the trade-off of heat and sucrose does not in itself represent new evidence about bee sentience, even when sucrose reward is predicted from colour rather than sensed directly. We already know that bee behaviour is guided by predicted future reward (7), as when they fly to a distant flower, rather than direct sensing of sucrose as in microbe chemotaxis. We also already know that bees can learn to associate reward with arbitrary stimuli when making these predictions (7).

Maybe the interaction between nociception and reward is what is critical? Figure 1C and 1D shows a simple situation in which the common value of a given heat/sugar combination is fixed, and bees make decisions between two options based on the differences between these fixed values. In this case, there would be no interaction between heat and sucrose concentration.

### Underlying signal detection theory

To explore the interaction further, we ran an analysis using signal detection theory.

In the original paper, the authors fitted their binary data with a logit link function. Their fixed-effects model was

“Proportion ∼ Concentration *Temperature”, where “Proportion” is the proportion of high-value choices, Temperature codes whether any feeders were heated or not (reference level = unheated), and “Concentration” is the sucrose concentration of the varying feeders (10%, 20%, 30%, 40%). It is helpful to think through the underlying signal-detection theoretic model implied by this. Effectively, we are modelling the expected difference in net value between the two choices, *Δ*, as sketched in Figure 1C and D.

This difference in expected value presumably declines as the concentration of the varying feeder approaches that of the high-value feeder, falling to zero when neither feeder is heated, and both contain 40% sucrose (blue line in Figure 2B). The difference in value is also reduced when the high-value feeder is heated (red line in Figure 2B is below the blue line). This explains why bees tend to avoid the heated feeder when both feeders offer 40% sucrose.

#### Interpretation of an interaction term

If there is a negative interaction between Temperature and Concentration, the decline would be steeper in the heated condition (red line steeper than blue line, as shown in Figure 2B). This would mean that bees treat heat as effectively less aversive when the relative reward is higher, potentially representing a motivational trade-off. Note that a positive interaction term does not make sense from this point of view, as it would mean bees treat heat as more aversive when there is more to be gained by enduring it.

Interestingly, an interaction term rules out the simple decision model sketched in Figure 1C and 1D. There, we assumed that each combination of sucrose S and heat H had a distinct value to the bee, say V(S,H). In Figure 1, we further represented this as linear, V(S,H) = S-H, but we can relax that assumption and write V as an arbitrary function of S and H. We assumed that the expected value of the difference between two choices was just the difference in these fixed values, *Δ* = V(S1,H1) - V(S2, H2). If this were valid, then for the bees’ decisions in the experiment we would have

*Δ* (Unheated,*C*) = V(40%,0) - V(*C*,0) ; bees choose between unheated feeders offering 40% or concentration *C*

*Δ* (Heated,*C*) = V(40%,*H*) - V(*C*,0) ; bees choose between a heated feeder offering 40% or an unheated feeder offering concentration *C*

By subtracting these, we can see that [*Δ*(Unheated,*C*)−*Δ*(Heated,*C*)] would be constant, equal to [V(40%,0)-V(40%,*H*)] regardless of the sucrose concentration *C* in the alternative feeder. This is consistent with a GLMM with no interaction term. In that case, the difference is equal to *β*_*H*_ (following terminology in Fig 2B). However, it is not consistent with an interaction term.

Thus, an interaction term means that we can’t model each option as having a fixed value to the bee, so that the difference between these options is just the difference in these fixed values. Rather, the relative value between the choices depends on the particular choices being made. Thus, a significant interaction term could be viewed as showing bees are making the sort of flexible motivational trade-off taken as evidence for sentience.

### Re-analysis of experimental data

The original paper claimed a marginally significant interaction term (p = 0.04, Figure 3A). This would imply that the aversive effect of a noxious stimulus is not fixed but depends on the relative value to be gained by enduring it. Some might consider this “enough flexibility to indicate centralized, integrative processing of information involving a common measure of value” (1). However, when we reanalysed the data, we found flaws that call this conclusion into question. We reanalysed all data in R Studio (2024.04.2).

**Figure 3.**
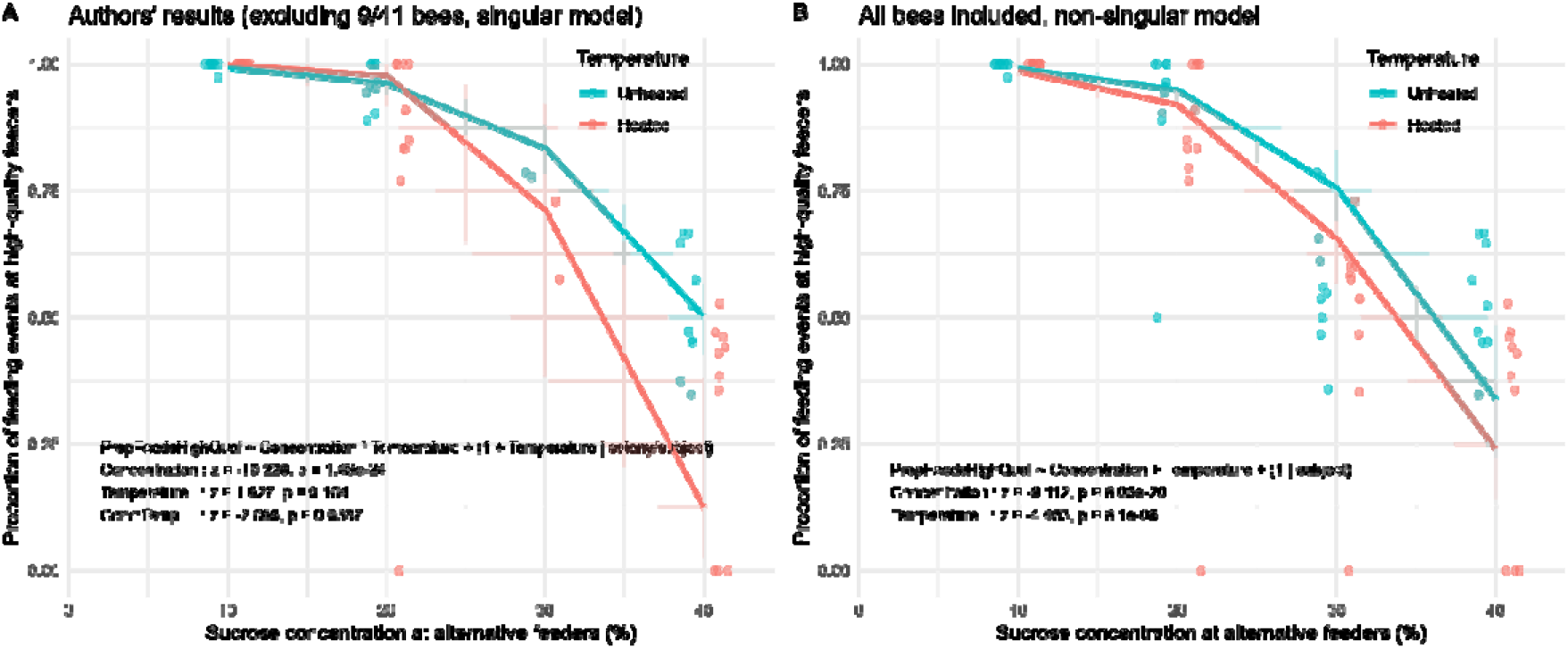
Authors’ analysis and our re-analysis. A: The authors’ included data and statistical model. 8/10 bees tested with 30% sucrose are excluded because, in the unheated condition, they did not show a significant preference for 40% over 30%; 1/10 bee in the 20% condition was also excluded. The model includes a complex set of random effects (1 +Temperature | colony / subject) and is singular. Statistical analysis finds no main effect of temperature (p=0.1) and a marginally significant interaction with concentration (p=0.04). B: Our re-analysis including all 41 bees tested and simpler random effects. We find no evidence for an interaction between temperature and concentration, but we do find highly significant main effects of both temperature and concentration (p<10^−5^). Dots show data for individual bees. There are two dots for each bee: heated and unheated. Lines show predictions of the fitted model, formula shown on the plot, using function glmer of R package lme4 (version 1.1-31). Note that the model in B is fitting more data with fewer parameters. Shaded regions show 95% confidence intervals obtained using function of bootMer of lme4 with 100 samples.

In the original paper, the authors excluded nearly a third (9 out of 31) of the bees tested in the crucial “trade-off” conditions comparing high and low sucrose because, during the initial unheated trials, these bees did not show a statistically significant preference for high sucrose. But bees were *not* excluded based on their performance in the heated conditions, or in the equal-sucrose condition. This is a problem because it could introduce bias. The paper’s conclusion requires that the proportion of high-value choices is lower in the heated condition than the unheated. Excluding bees that didn’t make enough high-value choices in the unheated condition, without applying such a criterion to the heated condition, risks printing an effect of temperature into the chosen data, whether or not it was there to begin with. Another result of excluding the data is that this leaves only two data points for the 30% condition.

There were also issues with the random effects used in the statistical model. In both the original paper and our reanalysis, models were fit with the package lme4 (version 1.1-31) using the command glmer. The code used in the paper fitted the formula

“Proportion ∼ Concentration *Temperature + (1+Temperature|colony/subject)”, where “Proportion” is the proportion of high-value choices, Temperature codes whether any feeders were heated or not (reference level = unheated), and “Concentration” is sucrose concentration of the varying feeders (10%, 20%, 30%, 40%). This formula fits random effects for subject nested within colony and also fits not only a random intercept but also a random slope (i.e. it allows the effect of heat to vary between bees). With only 2 results for each subject and <5 subjects per colony, there is not enough data to fit both random intercepts and slopes for Temperature within subject and colony, and so glmer warns that the model is singular. The results of this singular fit are shown in Figure 3A, which matches the results given in the paper (interaction: z=-2.068, P=0.039; main effect of Temperature : z = 1.627, P=0.104, n=32). The main effect of temperature is not significant, but the marginally significant negative interaction term indicates that bees become progressively less likely to select the heated feeder as the concentration in the unheated feeder approaches the high value of the heated feeder, consistent with a trade-off between desirable sucrose and undesirable heat.

To avoid the warning about singularity, we simplified the model to

“Proportion ∼ Concentration * Temperature + (1||)”

With the same bees excluded as in the paper, we now find a stronger interaction between Concentration and Temperature (z = 2.651, P= 0.008, n = 32), plus a significant main effect of Temperature (z= -3.727, P=0.0002, n=32). Since the reference temperature is Unheated, the negative main effect of Temperature indicates that bees are less likely to choose high-quality sucrose when these feeders are heated, as expected. However, the positive value of the interaction term means that the aversive effect of heat becomes actually becomes *less* as the concentration in the unheated feeder approaches the 40% available from the heated feeder, which is the opposite of what we would expect from a motivational trade-off.

The interaction term was driven entirely by one subject (bee ID 40) that was tested with 20% sucrose and never selected the 40%-sucrose feeder in the heated condition. With this bee removed, there was no significant interaction between Temperature and Concentration (interaction: z=-0.545, p = 0.586, n=31) although both had significant main effects when a model was fitted without an interaction term. This is true both for our model with simpler random effects, and for the original singular model: removing bee 40 abolished the interaction. Thus, the interaction term depends on excluding 9/41 bees, and even after that on just one of the remaining 32 bees. It is then significantly negative only when the model is singular due to overfitting the random effects and is significantly positive for a non-singular model where subject is the only random effect. This is thus not evidence for any negative interaction and therefore for a motivational trade-off.

If we run the “Concentration * Temperature + (1|subject)” model but including all 41 bees, again neither the interaction nor the main effect of Temperature are significant (interaction: z=-0.481, p = 0.631 ; main effect : z=-0.543, p = 0.587, n=41, AIC=473.6). If we drop the interaction term and fit “Proportion ∼ Concentration + Temperature + (1|subject)”, we obtain the model shown in Figure 3B (main effect of Temperature: z = -4.462, p = 8e-6, n=41, AIC=471.8).

We find main effects both of concentration and temperature, which are substantially more significant than the marginally significant interaction reported in the paper. However, we again find no evidence to support an interaction term, as required for a motivational trade-off.

## Discussion and conclusions

How to assess pain in animals is an important and fascinating question. Some suggested methods of assessing this have included investigating the impact of a noxious stimulus on future decision making and on analgesia seeking (8). Motivational trade-offs with pain have also been used as a key criterion. To rule out a simple behavioural trade-off, where reward and pain are combined in a fixed way, our analysis highlights the potential significance of a negative interaction between the response to the positive stimulus and to the painful stimulus. This has not always been taken into consideration. For example, when studying electric shock tolerance for hermit crabs in high- or low-quality shells, significant differences were found in the behaviour of crabs in high- or low-quality shells (3). However, the study did not statistically test for interaction effects between shell quality and pain tolerance. Recall that we’d similarly expect microbes to show higher tolerance for heat in the presence of higher sucrose concentrations if both of these stimuli control flagellar rotation by the same molecular pathway - but it seems a stretch to call this sentience.

Gibbons et al (5) pioneered a study that examined the question of motivational trade-offs in bees and crucially tested for interaction effects between sucrose preference and heat tolerance. This study allows us to begin to explore key questions on insect sentience and pain. However, our reanalysis of their data does not provide support for their conclusions.

The question of insect sentience is important both scientifically and for its implications. Given this, it is vital that we rely on strong evidence. We here argue that a significant negative interaction between responses to noxious and rewarding stimuli would imply a motivational trade-off and thus be suggestive of sentience, while significant main effects only are consistent with a purely behavioural trade-off such as found in the simplest biological systems. Ideally, neurophysiological evidence should help clarify if these trade-offs are made in the central nervous system and the mechanisms underlying them. Such studies will shed light on how a common value signal is neurally constructed and how flexible it really is. Recommendations for future studies would also include pre-registering the statistical analysis and any planned exclusion criteria. Criteria that introduce bias or result in the exclusion of a large number of subjects should be avoided. Certainly, any implications for policy should rely on robust designs and replications from multiple labs.

## Supporting information

Statistical Analysis

ReadMe File for Data Variable Names

## Acknowledgements

VN is funded by a BBSRC David Phillips fellowship BB/S009760/1 and a Leverhulme Trust Research Project Grant RPG-2021-358

## Data availability statement

Our code is provided as supplementary material. We thank the original authors for making the data and code available to us. These are uploaded at this link: https://doi.org/10.6084/m9.figshare.c.6066371.v4.

## Declaration of Interest

none

## Author’s contributions

VN and JR re-analysed the data and wrote the paper. JR developed the signal detection model.

## Ethical note

This article is based on a re-analysis of previous data and did not involve any experimentation on animals or humans.

## Notes

https://doi.org/10.6084/m9.figshare.c.6066371.v4

### Competing Interest Statement

The authors have declared no competing interest.

### Summary of Updates

The main text has been edited to streamline the reading, make the results more generally applicable and better acknowledge the contributions of the original paper.

