## Supplementary material for "Motivational trade-offs as evidence for sentience in bees: a critique": Statistical Analysis: MotivationalTradeoffsCritique2025_StatisticalAnalysis.R

```
rm(list=ls())
library(readxl)
library(ggplot2)
library(dplyr)

# Read in data file supplied by authors:
data = read_excel("MotivationalTradeoffsCritique2025_DataforRanalysis_DAPR.xlsx")

# Select only rows for individual trials:
data = data[grepl("^\\d+$", data$trial), ]

# make new columns for convenience:
data$NFeedsTotal =  as.integer(data$weightnormalaltupside)
data$PropFeedsHighQual =  data$propnormalaltupside
data$NFeedsHighQual =  as.integer(data$NFeedsTotal * data$PropFeedsHighQual)
# Where no feeds were made, we don't know the proportion of High Qual, so set that to NA
data$PropFeedsHighQual[data$NFeedsTotal==0]=NA

# Make a verbal label:
# In the origina data "Temperature" is a binary variable, 0 or 1
# I will call this "Heated":
data$Heated = data$Temperature 
# and will rewrite "Temperature" to be a string (easier for legends)
data$Temperature[data$Heated==0] = "Unheated"
data$Temperature[data$Heated==1] = "Heated"
# Make sure these are correctly considered factors
data$Temperature = factor(data$Temperature)
data$Temperature <- relevel(data$Temperature, ref = "Unheated")
data$subject = factor(data$subject)
data$colony = factor(data$colony)
# Let's also rename "Condition" with the more informative "Conc":
data$Concentration = data$Condition
data$Condition = NULL

##########################################################################
# Checking exclusions
##########################################################################

# NB the exclusion criteria the authors applied were:
# "Changed their 'excluded' variable to "no" (ie not excluded) 
# if they were in 10, 20 or 30 conditions and their result was significantly 
#different from 0.5, or if they were in 40 condition and their result was NOT significantly
#different from 0.5. The exclusion criteria applied to the unheated condition either, but the bee
# was excluded from analysis in both conditions (reasonably, the bias would have been even worse otherwise!)

# First of all we looked at the bees and checked we agreed which bees should be excluded on this basis.
# We did agree!
for (bee in unique(data$subject))
{
  unheatedtrials = dplyr::filter(data,subject==bee & Temperature=="Unheated")
  test = binom.test(x = sum(unheatedtrials$NFeedsHighQual),n = sum(unheatedtrials$NFeedsTotal), p=0.5)
  if (test$p.value>0.05 & mean(unheatedtrials$Concentration)<40)
  {
    print(sprintf("We find authors should exclude bee %s; marked as %s",bee,unheatedtrials$excluded[1]))
  }
}

# Remove excluded bees: - using authors' exclusion criterion
included = dplyr::filter(data,excluded!="yes")


# Was heat painful? Let's look at bees in the 40/40 condition
unheatedsameconc = dplyr::filter(data,Concentration==40 & Temperature=="Unheated")
nHigh = sum(unheatedsameconc$NFeedsHighQual)
nTotal = sum(unheatedsameconc$NFeedsTotal)
test = binom.test(x = nHigh,n = nTotal, p=0.5)
print(sprintf("On unheated same-conc (40%%/40%%) trials, %d out of %d feeds were towards the colour which would later be heated, p=%f",nHigh,nTotal,test$p.value))

heatedsameconc = dplyr::filter(data,Concentration==40 & Temperature=="Heated")
nHigh = sum(heatedsameconc$NFeedsHighQual)
nTotal = sum(heatedsameconc$NFeedsTotal)
test = binom.test(x = nHigh,n = nTotal, p=0.5)
print(sprintf("On heated same-conc (40%%/40%%) trials, %d out of %d feeds were towards the colour which was heated, p=%f",nHigh,nTotal,test$p.value))

# Which bees hate heat?
hateheat =  integer(0);
for (bee in unique(unheatedsameconc$subject))
{
  heatedtrials = dplyr::filter(data,Concentration==40 & subject==bee & Temperature=="Heated")
  nHigh = sum(heatedtrials$NFeedsHighQual)
  nTotal = sum(heatedtrials$NFeedsTotal)
  test = binom.test(x = nHigh ,n = nTotal, p=0.5)
  if (test$p.value<0.05 )
  {
    print(sprintf("Bee %s significantly avoided heat: Prop = %3.2f",bee,nHigh/nTotal))
    hateheat = c(hateheat,bee)
  }
}
NSbees = dplyr::filter(data,Concentration==40 & Temperature=="Heated" & !subject %in% hateheat)
nHigh = sum(NSbees$NFeedsHighQual)
nTotal = sum(NSbees$NFeedsTotal)
test = binom.test(x = nHigh,n = nTotal, p=0.5)
print(sprintf("For bees which individually showed NS effect of heat, on heated same-conc (40%%/40%%) trials, %d out of %d feeds were towards the colour which would later be heated, p=%f",nHigh,nTotal,test$p.value))


################################################################################
# Some useful functions;

#------------------------------------------------------------------------------
# Average all data for one bee over trials conducted in the same conditions:
avovertrials = function(data) {
  avdata = as.data.frame(data   %>% 
                               dplyr::group_by(subject,colony,Concentration,Temperature,Heated) %>% 
                               dplyr::summarize( NFeedsTotal=sum(NFeedsTotal),NFeedsHighQual=sum(NFeedsHighQual)))
  avdata$PropFeedsHighQual = avdata$NFeedsHighQual/avdata$NFeedsTotal
  return(avdata)  
}
#------------------------------------------------------------------------------
# Plot data as they do in the paper, including model predictions:
makeplotwithmodel = function(data,model,CI=95,nsim=100,title){
  
  # Make a dataframe for plotting model predictions:
  new_data <- expand.grid(
    Concentration = unique(data$Concentration),
    Temperature = unique(data$Temperature),
    colony = unique(data$colony)  # add a random colony or set to 1 for random effect
  )
  # Add predictions to the new data frame
  predictions <- predict(model, newdata = new_data, re.form = NA, type = "response")
  new_data$Predicted_PropFeedsHighQual <- predictions
  # Are we bootstrapping?
  if (nsim>0){
    # Get confidence intervals by bootstrapping (NBV this is slow, hence the low default for nsim)
    predict_function <- function(model) {
      predict(model, newdata = new_data, re.form = NA, type = "response")
    }
    boot_results <- lme4::bootMer(model, predict_function, nsim)
    new_data$CI_low <- apply(boot_results$t, 2, quantile, probs = (50-CI/2)/100)
    new_data$CI_high <- apply(boot_results$t, 2, quantile, probs = (50+CI/2)/100)
  }
  
  # Make a nice table to show on the plot
  # Extract z-values and p-values from the model summary
  model_summary <- summary(model)
  z_values <- model_summary$coefficients[, "z value"]
  p_values <- model_summary$coefficients[, "Pr(>|z|)"]
  
  # Format the text to display in the plot
  model_formula <- paste(deparse(formula(model)), collapse = "")
  model_formula <- gsub("\\s{2,}", " ", model_formula)
  
  # Figure out if the model includes an interaction term or not:
  if (is.na(z_values["Concentration:TemperatureHeated"])){
    annotation_text <- paste(
      model_formula,
      sprintf("Concentration : z = %5.3f, p = %4.3g", z_values["Concentration"], p_values["Concentration"]),
      sprintf("Temperature   : z = %5.3f, p = %4.3g", z_values["TemperatureHeated"], p_values["TemperatureHeated"]),
      sep = "\n"  )
  } else {
    annotation_text <- paste(
      model_formula,
      sprintf("Concentration : z = %5.3f, p = %4.3g", z_values["Concentration"], p_values["Concentration"]),
      sprintf("Temperature   : z = %5.3f, p = %4.3g", z_values["TemperatureHeated"], p_values["TemperatureHeated"]),
      sprintf("Conc:Temp     : z = %5.3f, p = %4.3g", z_values["Concentration:TemperatureHeated"], p_values["Concentration:TemperatureHeated"]),
      sep = "\n"  )
  }
  
  # Do the plots:
  g = ggplot() + 
    # Plot the data:
    geom_point(data=data,aes(x=Concentration,y=PropFeedsHighQual,group=Temperature,fill=Temperature,color=Temperature),position = position_jitterdodge(  jitter.width = 2, jitter.height = 0,dodge.width = 4, seed=1),alpha=0.6) +
    ggtitle(title) +
    # Set specific colors for "Heated" and "Unheated"
    scale_color_manual(values = c("Heated" = "#F8766D", "Unheated" = "#00BFC4")) +
    scale_fill_manual(values = c("Heated" = "#F8766D", "Unheated" = "#00BFC4")) +
    # Plus model predictions
    geom_line(data=new_data,aes(x = Concentration, y = Predicted_PropFeedsHighQual, color = Temperature, group = Temperature), size = 1) +  # line for each Temperature
    scale_y_continuous(limits = c(0, 1)) +  # Scale y-axis for binomial response
    theme_minimal() +
    theme(legend.position = c(0.8,0.9)) +
    annotate("text", x = 2, y = 0.1, label = annotation_text, hjust =0, vjust = 0, size = 3, color = "black") + 
    xlab("Sucrose concentration at alternative feeders (%)") +
    ylab("Proportion of feeding events at high-quality feeders")
  # Are we bootstrapping?
  if (nsim>0){
    g = g + geom_ribbon(data=new_data,aes(x = Concentration,ymin = CI_low, ymax = CI_high, fill = Temperature), alpha = 0.2)   # confidence interval ribbon
  }
  print(g)
  return(g)
}


###########################################
# Key results for paper
CI = 95 # We will show 95% confidence intervals on model predictions
nsim = 100 # We will get the confidence intervals by bootstrapping this many simulated values (it's really slow!)
print("#############################################################")
print("Let's pull together the key results")
print("#############################################################")
print("Point 1: We can recover the authors' original results with their analysis:")
# Using only data included in the paper
# For plotting purposes, let's average over all trials for a given bee/temp: 
# (NB as you would expect, you get exactly the same result with the model whether you set data=included or data=avincluded.
# But averaging is clearer for plotting)
avincluded = avovertrials(included)
# Using the authors' original model - NB this is singular because it attempts to fit both intercept & slope as random factors
# and to nest subject within colony. There is not enough data to support such a complex model

# The exact p- and z-values depend on the version or R and lmer used. Presumably this reflects the singularity of the model -
#  many different fits are equally good so what you get depends on random things

modelPaper = lme4::glmer(formula = PropFeedsHighQual ~ Concentration*Temperature + (1+Temperature|colony/subject),
                         weights = NFeedsTotal, family = "binomial",
                         data=avincluded)
print(summary(modelPaper))
g1=makeplotwithmodel(avincluded,modelPaper,CI,nsim,"Authors' results (excluding 9/41 bees, singular model)")


print("#############################################################")
print("Point 2: If we include all the data, there is a small but significant main effect of temperature:")
avdata = avovertrials(data)
modeldata = lme4::glmer(formula = PropFeedsHighQual ~ Concentration+Temperature + (1|subject),
            weights = NFeedsTotal, family = "binomial",
            data=avdata)
print(summary(modeldata ))
g2=makeplotwithmodel(avdata,modeldata,CI,nsim,"All bees included, non-singular model")


g=cowplot::plot_grid(g1, g2, labels = c('A', 'B'), label_size = 12)
ggsave(filename = "FigModel.tiff", plot = g, device = "tiff", 
       width = 12, height = 5, dpi = 300, units = "in", compression = "lzw")


###############################################
# Values reported in the statistics section:
print("The code used in the paper fitted the formula  proportion ~ Concentration *Temperature + (1+Temperature|colony/subject),  where proportion is the proportion of high-value choices, Temperature codes whether any feeders were heated or not, and “Concentration” is sucrose concentration of the varying feeders (10%, 20%, 30%, 40%). This formula fits random effects for subject nested within colony, and also fits not only a random intercept but also a random slope (i.e. it allows the effect of heat to vary between bees). With only 2 results for each subject and <5 subjects per colony, there is not enough data to fit both random intercepts and slopes for Temperature within subject and colony, and so glmer warns that the model is singular. ")
print(summary( lme4::glmer(formula = PropFeedsHighQual ~ Concentration*Temperature + (1+Temperature|colony/subject),
                               weights = NFeedsTotal, family = "binomial",
                               data=avincluded)))


print("To avoid this warning, we therefore simplified the model to  proportion ~ Concentration * Temperature + (1|subject). With the same bees excluded as in the paper, this actually strengthened the results. The paper reported a marginally significant interaction between Condition and Temperature (z = -2.068, P = 0.039, n = 32), and their model finds no significant main effect of Temperature (z=1.627, P=0.104, n=32, though this was not reported in the paper). With our simplified random effects, the model finds a stronger interaction between Condition and Temperature (z = - 2.651, P= 0.008, n = 32), plus a significant main effect of Temperature (z= -3.727, P=0.0002, n=32).")
print(summary( lme4::glmer(formula = PropFeedsHighQual ~ Concentration*Temperature + (1|subject),
                           weights = NFeedsTotal, family = "binomial",
                           data=avincluded)))


print("However, this was driven entirely by one subject, bee ID40, that was tested with 20% sucrose and never selected the 40%-sucrose feeder in the heated condition. With this bee removed, there was no significant interaction between Temperature and Concentration (interaction: z=-0.545, P=0.586, n=31) although both had significant main effects when a model was fitted without an interaction term. Thus, the interaction term depends entirely on excluding 9/41 bees, and even then rests solely on one of the remaining 32 bees. ")
print(summary( lme4::glmer(formula = PropFeedsHighQual ~ Concentration*Temperature + (1|subject),
                           weights = NFeedsTotal, family = "binomial",
                           data=dplyr::filter(avincluded,subject!=40))))


print("If we run the same model but including all 41 bees, neither the interaction nor the main effect of Temperature are significant (interaction: z=-0.543, P=0.587; main effect : z=-0.481, P=0.631, n=41). ")
print(summary( lme4::glmer(formula = PropFeedsHighQual ~ Concentration*Temperature + (1|subject),
                           weights = NFeedsTotal, family = "binomial",
                           data=avdata)))

print("If we run the same model but including all 41 bees, neither the interaction nor the main effect of Temperature are significant (interaction: z=-0.543, P=0.587; main effect : z=-0.481, P=0.631, n=41, AIC=473.6). If we drop the interaction term and fit proportion ~ Concentration + Temperature + (1|subject) , we obtain the model shown in Figure 1B (main effect of Temperature:  z = 4.462, P=8e-6, n=41, AIC=471.8).")
print(summary( lme4::glmer(formula = PropFeedsHighQual ~ Concentration + Temperature + (1|subject),
                           weights = NFeedsTotal, family = "binomial",
                           data=avdata)))
```
