## Supplementary material for "Motivational trade-offs as evidence for sentience in bees: a critique": ReadMe File for Data Variable Names

Readme file for the data uploaded along with **Motivational trade-offs as evidence for sentience in bees: a critique**

Please note that the this is based on data provided by the authors of the target article. Since that datafile does not have a readme, the details below are based on their code and data file. We only provide the details of the variables used in our analysis.

1. Trial: The trial number
2. Weightnormalaltupside: Total number of feeds
3. Propnormalaltupside: Proportion of feeds at the high quality feeder
4. Temperature: Whether the feeded was heated or not, binary variable.
5. Subject: Identity of the bee
6. Colony: Colony identity of the bee
7. Condition: Concentration of the alternative feeder
